# Autophagy is dispensable in germline stem cells but is required in the cap cells for their maintenance in the Drosophila ovarian niche

**DOI:** 10.1101/2024.10.21.619512

**Authors:** Kiran Suhas Nilangekar, Bhupendra V. Shravage

## Abstract

Autophagy is a cytoprotective mechanism responsible for the maintenance and long-term survival of various cell types, including stem cells. However, its role in the Germline stem cell (GSC)-niche, including its effects on GSCs and niche cells, remains unexplored. We demonstrate that autophagy flux in female Drosophila GSCs is low and dependent on the core autophagy gene, *Atg5*. However, the maintenance of *Atg5-/-* GSCs within the GSC-niche was unaffected. In contrast, disruption of autophagy within the cap cells (niche cells) leads to the loss of both cap cells and GSCs during aging. Further, reduced autophagy in cap cells severely impairs the crucial GSC self-renewal signal mediated by BMP-pMad emanating from the cap cells post-midlife. Autophagy was essential for the long-term survival of cap cells. Our study reveals a differential role for autophagy, which is dispensable in GSCs but necessary in niche cells, where it supports signaling and survival to maintain GSCs.

## Introduction

Macroautophagy (autophagy hereafter) is a catabolic process that maintains cellular homeostasis by the sequestration and lysosome-mediated degradation of cytoplasmic material that is toxic or superfluous. Autophagy occurs constitutively at the basal level and can be induced to elevated levels under stress conditions, including nutrient deprivation. Therefore, basal autophagy acts as a housekeeping mechanism, and autophagy is elevated in response to stress. The role of autophagy is particularly important and recognized in aging and longevity. Stem cell niches replace dysfunctional cells within the tissues and organs throughout the lifespan. However, the role of autophagy in the maintenance of stem cell niches throughout the lifespan is poorly understood. The stem cell niche consists of stem cells, ECM, signaling factors that regulate the stem cells, and non-stem cells (niche cells) that regulate stem cells, the stromal component, i.e., blood vessels, nerves, and the extracellular matrix. The niche maintains the “stemness” of stem cells by activating pro-stemness cellular pathways and repressing differentiation. Stem cells are also subject to cellular aging and have reduced proliferative capacity with increased age. The germline stem cell (GSC)-niche is well-studied in Drosophila and is a good model to address the role of autophagy in stem cell-niches.

The process of macroautophagy (autophagy hereafter) has been well characterized in invertebrate models like *C. elegans, Drosophila,* and mammals. Several Atg (autophagy-related) proteins govern this process. Autophagy is influenced by many pathways, including those that regulate cellular metabolism. Tor/MTORC1/mTOR (mechanistic target of rapamycin) is one of the central intracellular pathways that directly regulates autophagy. In Drosophila, mTOR kinase is active under nutrient-rich conditions and inactivates Atg1 and Atg13 through specific phosphorylation on serine/threonine residues. Under stress conditions, including nutrient deprivation, mTOR dissociates from the Atg1-Atg13 complex. Atg1, Atg13, Atg101, along with FIP200, form the induction complex called Atg1 complex. A second complex (called Vps34 complex) consisting of Atg14, Vps34, Atg6, and Vps15 catalyzes membrane nucleation. Atg18 proteins and Atg2 bind to the PI3K complex, resulting in a phagophore assembly site. Atg2 and Atg9 transport lipids/phospholipids required to expand the phagophore membrane through their lipid translocase and scramblase activities respectively. Two ubiquitin-like conjugation systems are essential for phagophore expansion. The first is the Atg12-Atg5-Atg16 conjugation system. In this system, Atg12 covalently binds to Atg5 through the action of E1-like activating enzyme Atg7 and E2-like conjugation enzyme Atg10. The Atg12-Atg5 complex interacts with Atg16 to form the Atg12-Atg5-Atg16 complex that functions as an E3-like ligase for Atg8, a ubiquitin-like component of the Atg8 conjugation system. Atg4 cleaves the C-terminal residue on Atg8 in this system to expose the glycine. The E1-like activating enzyme Atg7 further activates cleaved Atg8 (Atg8-I) and transfers it to the E2-like enzyme Atg3. Atg12-Atg5-Atg16 complex catalyzes the covalent conjugation of Atg8-I with the lipid phosphatidylethanolamine (PE) present on the phagophore for form Atg8-II/Atg8-PE, a process is called ‘lipidation of Atg8. ‘Autophagy receptors’, including Ref(2)P and Nemo, can act as adaptors by bridging cargo and Atg8, resulting in active and selective cargo sequestration. Repeated actions of these Atg protein complexes complete phagophores to form double-membraned sacs/vesicles called “autophagosomes” with the cargo inside. Autophagosomes are transported to lysosomes or vice-versa via molecular motors interacting with LC3 along microtubules. The fusion of autophagosome and lysosome is mediated by SNAREs, particularly Syntaxin-17, SNAP29, and VAM7/9/VAMP8. The outer membrane of the autophagosome fuses with the lysosomal membrane to form ‘autolysosomes.’ The contents, along with the inner membrane of the autophagosomes, are degraded within the lysosomes by acid hydrolases. Lysosomal permeases and other channels present on the membrane mediate the efflux of the molecules produced by autolysosomal digestion [2,3].

Autophagy-related protein 5 (Atg5) is a core autophagy protein and an essential component of the Atg12-Atg5-Atg16 complex. The Atg12-Atg5-Atg16 complex is responsible for the lipidation of Atg8, without which the phagophore fails to expand and mature, indicating that Atg5 is indispensable for autophagy. Hence, knocking down or knocking out core autophagy genes, including *Atg5,* is a classical approach to studying autophagy [4]. Atg5 can also be used to mark immature autophagosomes in certain cellular contexts [5,6]. *Atg5* deficient mice do not survive post-natally [7]. In the context of the germline, ATG5 was shown to be required for sperm development and fertility in mice [8]. Importantly, ubiquitous overexpression of Atg5 in mice was sufficient to enhance the levels of autophagy and extend the lifespan [9]. Neural stem cells in mice with cell-type specific conditional knockout of *Atg5* showed impaired autophagy as observed by reduced LC3 lipidation and accumulation of p62 [10]. Importantly, these *Atg5* knockout NSCs had an accumulation of dysfunctional mitochondria. Additionally, mutations in *ATG5* were found to cause ataxia in humans [11]. In Drosophila, autophagy is disrupted in Atg5 knockdown mediated by *Atg5RNAi* and *Atg5* null mutants. Recent studies have reported instances of LC3 lipidation-independent autophagy, although these primarily represent forms of selective autophagy [12]. Given its essentiality for canonical autophagy, Atg5 is an ideal candidate for investigating autophagy and its role in the GSC-niche.

Autophagy is crucial adult stem cells and has been studied in the context of associated diseases, metabolism, stem cell self-renewal, and differentiation [13,14]. The role of autophagy for stem cell differentiation has been found to be crucial in multiple model organisms. The role of autophagy in stem cell aging has been well-studied in muscle stem cells [15] and hematopoietic stem cells (HSCs)[16]. For both systems, autophagy was demonstrated to be important in maintaining quiescence and regenerative function in old stem cells. Autophagy is defective in aged muscle stem cells in mice and humans. This defective autophagy in old stem cells and experimentally impaired autophagy in young stem cells is responsible for their entry into senescence. Due to reduced proteostasis, mitochondrial dysfunction and oxidative stress was elevated in these cells, which further resulted in reduced function and number of muscle stem cells. Studies have demonstrated that restoring autophagy in old stem cells reverses senescence and restores their regenerative functions. Interestingly, about one-third of the old hematopoietic stem cell population exhibits high levels of autophagy, low metabolism, and robust regenerative capacity [16]. These autophagy-activated old HSCs were fitter than the remaining low-autophagy old HSCs.

If autophagy is required to maintain niche cells and thereby indirectly influences GSCs is not clear. Our group has previously reported a role for Atg1 in GSC maintenance, but the mechanism is via regulation of mitochondrial dynamics and not necessarily as a function of autophagy [17]. Moreover, there has not been any direct investigation into the role of autophagy in the maintenance of niche cells. In Drosophila, researchers found that depletion of autophagy in male cyst stem cells (CySCs; part of niche) negatively affects their maintenance and leads to loss of GSCs. Previous reports also suggest that there is an age-dependent loss of niche cells, subsequently affecting stem cell maintenance. Aging has been shown to alter the secretome and transcriptome of the HSC-niche and neurogenic niche cells in mice [20,21]. The niche-secreted factors for spermatogonia stem cell/GSC self-renewal in male mice alter with age [23]. The niche-stem cell signaling communication axes were perturbed due to the aging of muscle stem cells [24]. Epidermal Growth Factor Receptor (EGFR) signaling responsible for the regulation of murine neural stem cells is reduced in the subventricular zone (niche) due to aging [25]. Thus, niche cells are responsible for maintaining stem cells and the deterioration of niche cells due to factors that disrupt homeostasis, including aging, leads to the loss of stem cells (reviewed in [19, 22]). Currently, there is no fundamental understanding of the molecular role of autophagy in any of the niche cells that comprise adult stem cell niches.

Drosophila has served as an amenable model for the study of autophagy [2,26]. Several studies in Drosophila have demonstrated the importance of autophagy during development, morphogenesis, cell death, nutrient stress and aging [27,28]. The female Drosophila GSC niche has been an excellent model for studying stem cell biology and stem cell-niche interaction. Stem cell niche was first described in female Drosophila and is a model for studying GSC aging [29] [30]. The female GSC-niche is present at the anterior tip of a structure called “germarium”. Germarium is the anterior-most part of the ovariole, a functional unit of the ovary, that harbors progressive developing eggs towards the posterior. The niche is made up of terminal filament cells, cap cells and escort cells. The primary niche cells, cap cells, secrete the Bone Morphogenetic Proteins (BMPs) pathway ligands Decapentaplegic (Dpp) and Glass bottom boat necessary for GSC self-renewal, and they also provide physical anchorage to the GSCs via adherens junction proteins including E-Cadherin. On an average there are six-eight cap cells and two-three GSCs in the GSC-niche. GSCs are maintained in the undifferentiated state by activation of the BMP-pMad signaling in the GSCs. Active BMP signaling, leads to the phosphorylation of Mothers against Dpp (Mad) in GSCs where it dimerizes with Medea and translocates into the nucleus to repress differentiation factors including bag-of-marbles (Bam).

Previous studies demonstrate an age-dependent reduction in the number and function of GSCs and cap cells [31]. The authors also found that the BMP/Dpp signaling and E-Cadherin mediated anchorage between the niche and GSCs decrease in older germaria. In addition to the intrinsic aging of stem cells, the age-dependent deterioration of niche cells was confirmed to be the extrinsic factor causing GSC aging. Similar to female GSCs, in the male Drosophila GSC-niche, there is an age-dependent reduction in the number of niche/hub cells, number of GSCs, E-Cadherin and the niche signaling factor Unpaired [32,33].

In this study, we determine whether autophagy is required intrinsically in the GSCs or extrinsically in the niche cells for stem cell maintenance during aging using female GSC-niche.

## Results

### Autophagy in GSCs is low and dispensable for GSC maintenance

Previously, we generated and characterized germline-specific autophagy reporter *nosP-mCherry-Atg8a*, which expresses mCherry tagged Atg8a under the *nanos* promoter [34]. Under normal nutritional conditions, we observed very few mCherry-Atg8a puncta in the female GSCs compared to the differentiating cysts in regions 2a and 2b of the germarium. Autophagy is strongly induced in response to nutrient limitation in the germarium [34–37]. However, we did not observe any change in mCherry-Atg8a puncta in the GSCs upon starvation, suggesting that GSCs may be protected from starvation-induced autophagy. Also, the mCherry-Atg8a puncta in GSCs did not change upon pharmacological treatments that upregulate or disrupt autophagy, specifically in response to exposure to rapamycin and chloroquine, respectively. In the present study, we performed a thorough quantitative measurement of autophagy flux in the GSCs. Flies expressing mCherry-Atg8a (marks autophagosomes and autolysosomes) were subjected to chloroquine treatment and immunostained for Cathepsin L (marks lysosomes and autolysosomes). Upon quantification, it was found that neither the number nor the size of mCherry-Atg8a puncta changed upon chloroquine treatment (Figure 1B and C). Also, the number of Cathepsin L puncta was low and did not change upon chloroquine treatment (Figure 1B and C). Importantly, on average, one punctum per GSC was positive for both mCherry-Atg8a and Cathepsin L, which denotes autolysosome, which also did not change upon chloroquine treatment (Figure 1B). In contrast, mCherry-Atg8a puncta accumulate upon chloroquine treatment in the differentiating cells in region 2 of the germarium [34]. These data confirm that basal autophagy and autophagy flux in GSCs is low.

**Figure 1:**
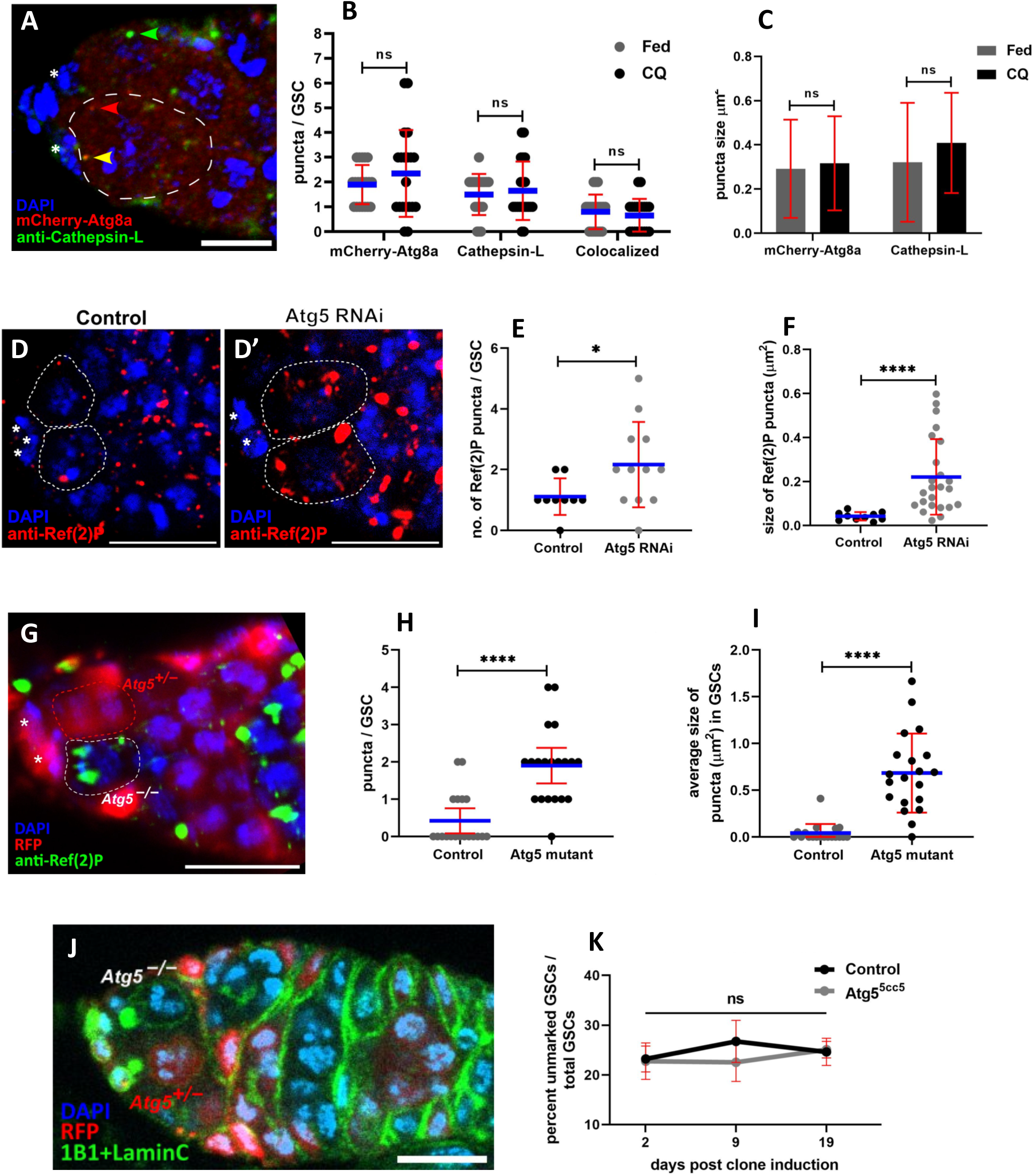
Autophagy in GSCs is low & dispensable for maintenance. (A) Representative image showing autophagic vesicles in GSCs. The GSC is marked by dashed outline and the cap cells are marked by asterisks. Puncta are indicated by colored arrowheads; mCherry-Atg8a/autophagosome (red), Cathepsin L/lysosome (green) and colocalized punctum/autophagosome (yellow). Scale bar 5 μm. (B) Interleaved scatter plot showing no difference in the number of autophagic vesicles in control GSCs and CQ-treated GSCs. Blue line represents the average and error bars represent standard deviation. (C) Bar graph showing no difference in the size of mCherry-Atg8a or Cathepsin L puncta in control GSCs and CQ-treated GSCs. n=20 GSCs per treatment. (D, D’) Representative image showing accumulation of Ref(2)P in the GSCs upon *Atg5* knockdown. The GSCs are marked by dotted outlines and the cap cells are marked by asterisks. Scale bar 10 μm. Interleaved scatter plots showing increase in (E) number and (F) size of Ref(2)P puncta in GSCs upon *Atg5* knockdown. Blue line represents average and error bars represent standard deviation. n=9 & 12 GSCs for control & Atg5 KD respectively. (G) Representative image showing severe accumulation of Ref(2)P in Atg5 null GSCs. Atg5 null (-/-) GSC is marked by white dashed outline (RFP–), heterozygous (+/-) GSC is marked by red dashed outline (RFP+) and cap cells are marked by asterisks. Scale bar 10 μm. Interleaved scatter plot showing increase in (H) number and (I) size of Ref(2)P puncta in Atg5 mutant GSCs. Blue line represents average and error bars represent standard deviation. Two independent experiments were performed. For the data presented, n=19 control GSCs which were heterozygous(+/-) or wildtype (+/+) & n=20 Atg5 mutant GSCs. (J) Representative image showing mosaic GSC clones. Atg5 null GSC (indicated as *Atg5*–/–) with no visible RFP and neighboring RFP positive heterozygous GSC (indicated as *Atg5*+/–). Scale bar 10 μm. (K) Line graph showing no difference in the maintenance of Atg5 null GSCs (Atg5^5cc5^) compared to control GSC clones. Error bars represent standard deviation. The data presented is cumulative from three independent biological replicates. n=105±6 (I), 80±2 (II) & 70±3 (III) germaria per genotype per time point for the three replicates (I, II and III). (except III Atg5^5cc5^ 2d n=54). \**p*<0.05, \*\*\*\**p*<0.0001.

Additionally, we measured autophagy flux under conditions where autophagy was induced genetically. Overexpression of core autophagy genes like *Atg1*, *Atg8a* or *Atg5* has been demonstrated to elevate autophagy flux [38,39]. Previous studies demonstrated that Atg5 overexpression can increase autophagic activity [9]. *Atg5* was overexpressed in the germline, including the GSCs, using *UASp-eGFP-Atg5* transgene driven by *nosGal4VP16* germline-specific Gal4 driver. eGFP fluorescence was visible in the germline cells, including the GSCs (Figure S1B). We quantified autophagy flux in the GSCs to test if *Atg5* overexpression was sufficient to increase the autophagy flux in GSCs in the presence and absence of CQ. Despite *Atg5* overexpression, we did not observe increased autophagy flux in GSCs, which was comparable to that in control GSCs (Figure S1A).

As recommended by Klionsky et al., 2021, autophagy should be assessed by two independent markers [4]. The autophagy receptor protein Ref(2)P was used as a marker to measure autophagic activity. Ref(2)P/p62/SQSTM1 is a protein that sequesters cargo for autophagic degradation and, in the process, itself gets degraded along with the cargo. Thus, typically the amount of Ref(2)P is inversely proportional to the autophagic activity. Also, Ref(2)P, if not degraded via autophagy, tends to form aggregates that can vary in number and size. We observed that there was no difference in the size and the number of Ref(2)P aggregates in the control GSCs and Atg5 overexpressing GSCs (Figure S1C and D). These data confirm that the autophagic activity in GSCs does not increase despite *Atg5* overexpression.

We next tested if Atg5 is necessary for maintaining basal autophagy within the GSCs. *Atg5* mRNA was depleted specifically in the germline by expressing *Atg5IR* using *nosGal4VP16* [34]. There was a significant increase in the number as well as the size of Ref(2)P puncta in GSCs upon *Atg5* knockdown as compared to control GSCs (Figure 1D, E, and F). Additionally, we used a null mutation of Atg5, *Atg5^5cc5^*, to confirm its requirement for basal autophagy. The *Atg5^5cc5^* mutation has a deletion of more than 85% of the coding region, including the translation start site; it does not code for any protein and hence is a null mutant [11]. We employed the FLP-FRT technique to generate Atg5-/- GSC clones. We observed a significant increase in the number as well as size of Ref(2)P puncta in Atg5 mutant (Atg5-/-) GSCs as compared to heterozygous (Atg5-/+) or wildtype (Atg5+/+) GSCs (Figure 1H and I). In summary, these data confirm that Atg5 is required for Ref(2)P clearance in the GSCs, indicating that canonical Atg5-dependent autophagy occurs in the GSCs.

Our data suggest that Atg5 is required for basal autophagy within the GSCs and previous studies have demonstrated that autophagy is crucial for stem cell maintenance. Hence, we examined the role of autophagy in the maintenance of GSCs using Atg5 null mutants. We performed a GSC retention assay, in which the rate of FLP-FRT-induced *Atg5* mutant (Atg5-/-) GSC clones retained was compared with those of control GSC clones (Atg5-/+ or Atg5+/+) (Figure 1J and K). Surprisingly, there was no difference in the retention of *Atg5* mutant GSCs compared to that of control GSCs even up to 19 days post-clone induction (Figure 1K). Therefore, Atg5-dependent autophagy is dispensable for GSC maintenance. Atg5 is necessary for autophagy in the GSCs but not for their maintenance, suggesting that autophagy appears to be dispensable for GSC maintenance.

### Autophagy is required extrinsically in the cap cells for GSC maintenance

In the Drosophila male GSC-niche, Sênos Demarco and colleagues observed that knocking down autophagy genes in the CySCs (cyst stem cells), which are part of the niche, significantly affects the GSC number [18]. In a similar approach, we wanted to establish and validate a model for the ‘autophagy-defective niche.’ We targeted the primary niche; cap cells and used the UAS-Gal4 enabled RNAi-mediated knockdown of the following Atgs: *Atg1, Atg13, Atg9, Atg5, Atg12, Atg16,* and *Atg8a*. The *Atg-*RNAi-mediated knockdown was achieved using cap cell-specific Gal4 drivers, HhGal4 or Bab1Gal4. We confirmed the disruption of autophagy by monitoring Ref(2)P aggregation in the *Atg*-knockdowns. Compared to control cap cells, we found a significant accumulation of Ref(2)P in all but two *Atg*-RNAi tested (Atg12 and Atg13)(Figure 2A, C and D, Figure S1E—H). Additionally, for *Atg5 RNAi*, we used the 3xmCherry-Atg8a reporter, which expresses the *mCherry-Atg8a* transgene under the control of the endogenous promoter. Cap cells with Atg5 knockdown had drastically reduced mCherry-Atg8a puncta as compared to controls (Figure 2B and E). Taken together, the autophagy-defective niche model could be established and validated for *Atg1, Atg9, Atg5, Atg16, and Atg8a*.

**Figure 2:**
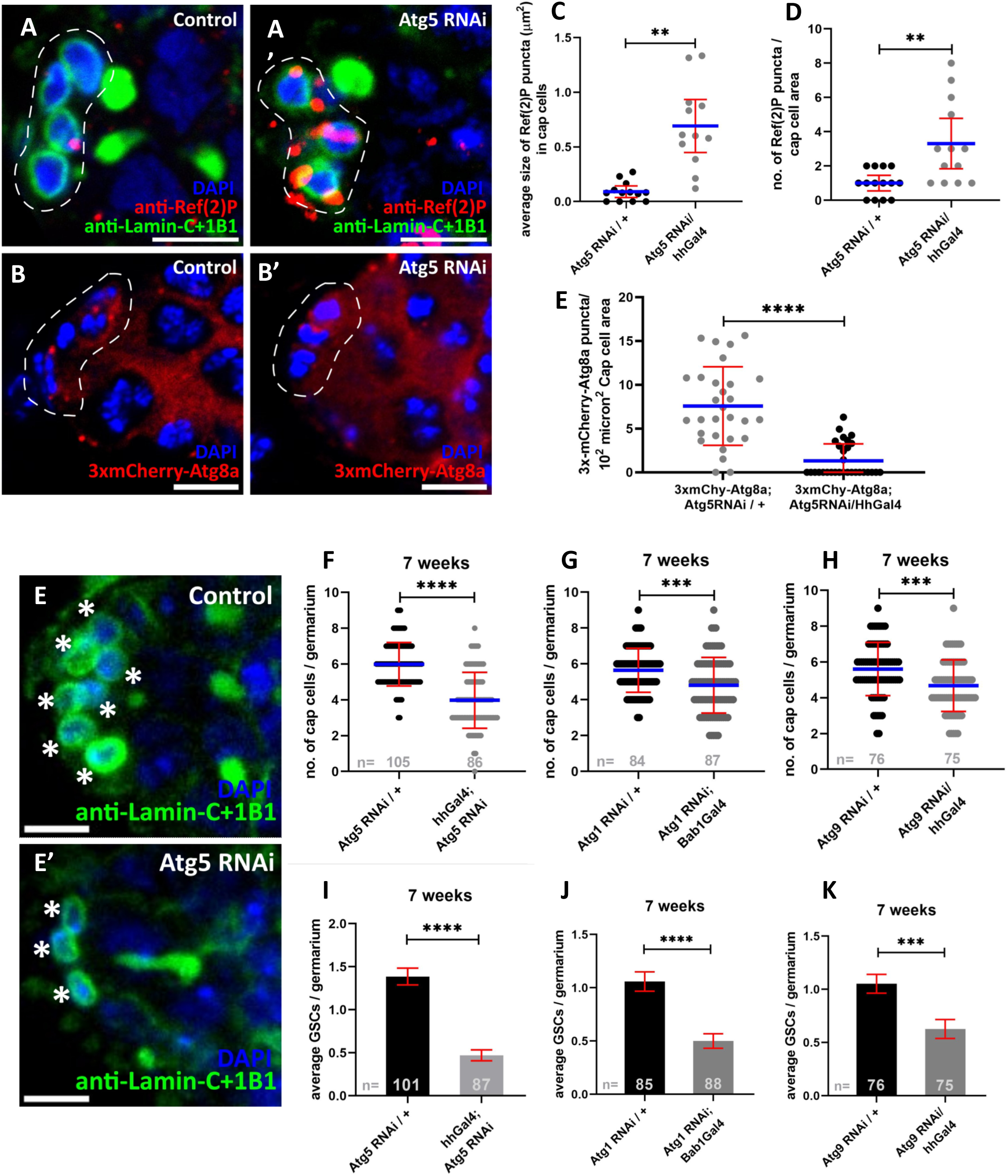
Autophagy in Cap cells is necessary for their and GSC maintenance. (A, A’) Representative image showing severe accumulation of Ref(2)P in cap cells. (B, B’) Representative image showing the reduction of autophagosomes in Atg5 RNAi cap cells. The region of cap cells is marked by dashed outlines. Scale bar 5 μm. Interleaved scatter plots showing increase in size (C) and number (D) of Ref(2)P puncta in cap cells with *Atg5* knockdown. n=14±1 cap cell planes/area per genotype. (E) Interleaved scatter plot showing the reduction of 3xmCherry-Atg8a puncta (autophagosomes) in Atg5 KD cap cells. n=30 cap cell planes/area per genotype. Blue line represents the average and error bars represent standard deviation. (E-E’) Representative image showing the loss of cap cells due to Atg5 KD. Cap cells are marked by asterisks. Scale bar 5 μm. Interleaved scatter plots showing the reduced number of cap cells in old niches with (F) Atg5 RNAi, (G) Atg1 RNAi and (H) Atg9 RNAi in the cap cells. Blue line represents the average and error bars represent standard deviation. Bar graphs showing the reduced number of GSCs in old niches with (I) Atg5 RNAi, (J) Atg1 RNAi and (K) Atg9 RNAi in the cap cells. Error bars represent standard error of mean. Sample sizes are the number of germaria as indicated in the graphs. \*\**p*<0.01, \*\*\**p*<0.001, \*\*\*\**p*<0.0001.

Previous studies have demonstrated that cap cells are essential for maintaining GSCs. We tested if autophagy in cap cells is essential for maintaining GSCs in a non-autonomous manner. To address this, we used the autophagy defective niche model of *Atg1*, *Atg9*, and *Atg5* knockdown and counted the number of GSCs retained in germaria of seven-week-old flies. Remarkably, there was a significant loss of GSCs in niches with *Atg1*, *Atg9* or *Atg5* knockdown in cap cells compared to age-matched controls (Figure 2I, J and K). Hence, autophagy is required in the cap cells for GSC maintenance during aging. Previous reports have demonstrated that the number of GSCs, is proportional to the number of cap cells (One GSC for about 2.5 cap cells [29][40]. Since we observed fewer GSCs in aged autophagy-defective niches compared to age-matched control, we quantified the number of cap cells in such GSC-niches. Notably, it was found that there were significantly fewer cap cells in autophagy-defective niches than in age-matched controls (Figure 2E, F, G and H). Since multiple Atgs were targeted, and all show the same phenotype, the rapid loss of cells in the GSC-niche during aging is attributable to the disruption of autophagy in the cap cells. Thus, autophagy is essential for maintaining cap cells, which in turn is necessary for the maintenance of GSCs (GSCs).

Since the HhGal4 and Bab1Gal4 drivers are active during development and gonad morphogenesis, it was critical to test whether the observed effects preceded the adult ovary stage. Therefore, in another experiment, cap cells and GSCs were quantified earlier, i.e., 12 days after eclosion. The average number of cap cells and GSCs in all the experimental conditions was found to be comparable to their respective controls at 12 days (Figure S2B, C and Figure 3G). These data suggest that the ratio of the number of cap cells to that of GSCs was similar in controls and Atg5 knockdown niche, confirming that morphogenesis of the GSC-niche was unaffected during development. Thus, the exacerbated loss of cells within the niche was due to the lack of autophagy during aging.

**Figure 3:**
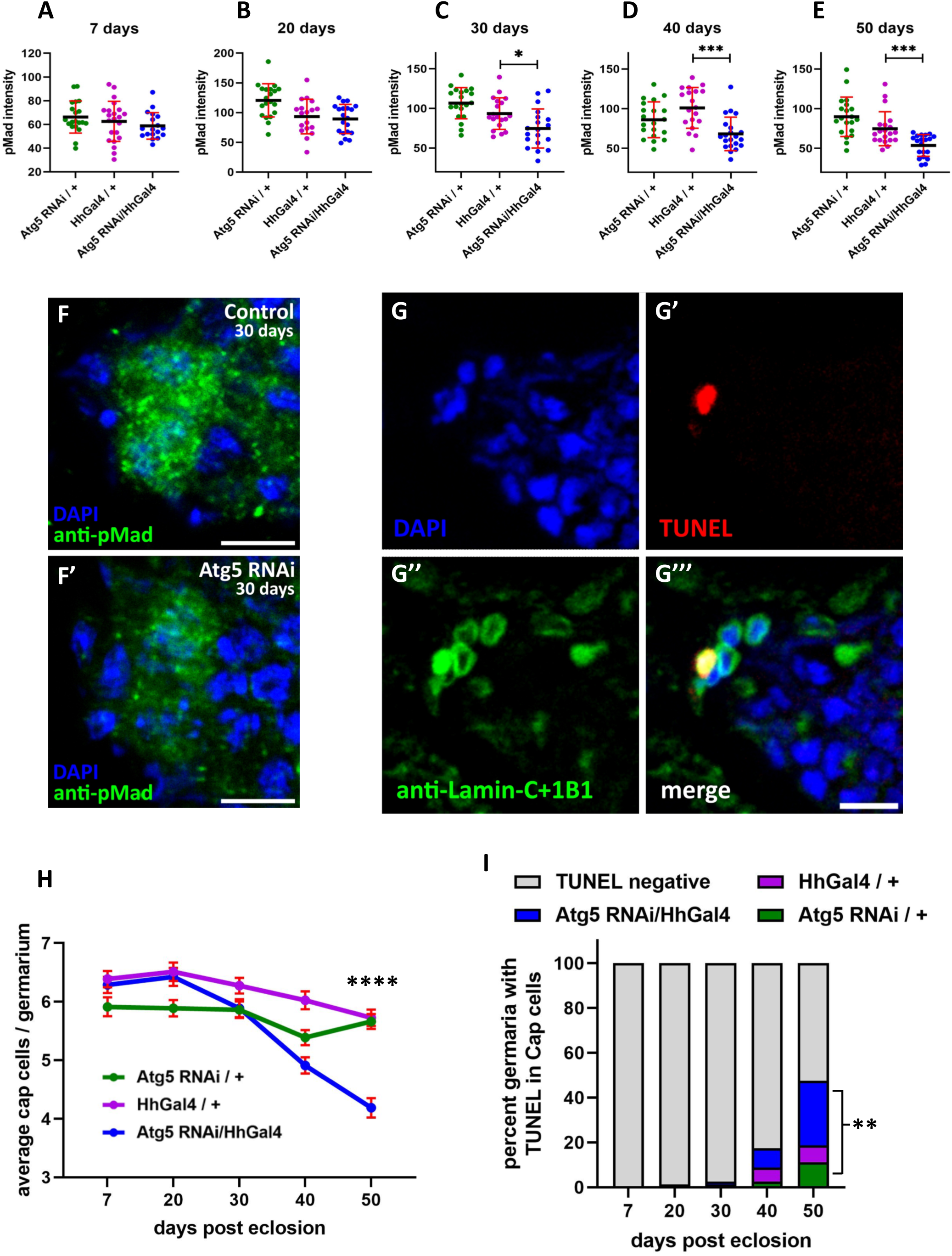
Lack of autophagy affects cap cell function and causes cap cell death during aging. (A—E) Interleaved scatter plots showing pMad intensity in GSCs of the mentioned genotypes at the five mentioned timepoints. Error bars represent standard deviation. n=20±2 GSCs per genotype per time point. \**p*<0.05, \*\*\**p*<0.001. (F, F’) Representative image showing reduced pMad in GSCs at 30 days due to *Atg5* knockdown in the cap cells. Scale bar 5 μm. (G—G’’’) Representative image showing TUNEL positive cap cell in the GSC niche. Scale bar 5 μm. (H) Line graph showing decline of cap cells due to Atg5 knockdown. Error bars represent standard error of mean. (I) Stacked column graph showing age-wise distribution of TUNEL positive cap cells out of total cap cells in the mentioned genotypes. n=80 germaria per genotype per time point.

### Lack of autophagy affects cap cell function and causes cap cell death during aging

In female Drosophila, GSC self-renewal is controlled by several factors, which include secreted and membrane-bound factors. One of the primary functions of cap cells is to secrete BMP ligands, Decapentaplegic and Glass bottom boats, which provide self-renewal signals to the GSCs and control their division. This function of secretion of BMPs by the cap cells is indispensable for GSC maintenance since studies have shown that the loss of BMP signaling leads to the activation of the stem cell differentiation program, leading to GSC loss [41]. Antibodies that specifically recognize phosphorylated Mad (pMad), a core signal transducer of BMP/Dpp signaling, have been extensively used as a reliable marker to quantify the strength of cap cell-GSC signaling. We quantified the pMad levels within GSCs in autophagy-deficient niches. We found a significant reduction in pMad intensity in GSCs from aged autophagy-defective niches 30 days onwards compared to those from age-matched controls (Figure 3A—F). Additionally, the same phenotype could be replicated using another cap cell-specific driver line, Bab1Gal4 (Figure S2D). Thus, impaired autophagy in the cap cells affected BMP-pMad signaling in the GSC-niche in middle-aged flies and later during aging.

In the autophagy-defective niches, we observed rapid loss of cap cells during aging (Figure 3I). Hence, we tested if the loss of cap cells was due to apoptosis. We performed the TUNEL assay, which facilitates identifying and quantifying cells undergoing apoptosis by fluorescently marking the sites of double-stranded DNA breaks that occur during programmed cell death. Double-stranded DNA breaks in control and autophagy defective niches during the lifespan were assayed using TUNEL staining, and the cap cells positive for TUNEL staining were quantified. Cap cell death was not observed in any of the experimental conditions until the 30-day time point. TUNEL-positive cap cells were present in control as well as autophagy-defective niches at 40-day timepoint. However, there were significantly more dying cap cells at 50 days, in *Atg5* knocked down, compared to age-matched controls (Figure 3 H and J). This suggests that lack of autophagy led to apoptosis in aged cap cells, indicating that autophagy functions as a survival factor within cap cells during aging.

## Discussion

In this study we have discovered a differential role of autophagy in the Drosophila ovarian GSC-niche. We demonstrate that autophagy is dispensable intrinsically within the GSCs, whereas it is required in the niche cells as an extrinsic factor for GSC maintenance. First, we quantified the basal autophagy flux in the GSCs and found that it was low and did not increase despite *Atg5* overexpression. Although *Atg5* was required for the clearance of autophagy cargo receptor Ref(2)P within the GSCs, the maintenance of *Atg5* null mutant GSCs was unaffected by the complete lack of autophagy. *Atg5*-mediated autophagy was required for Ref(2)P clearance within the cap cells and, in contrast to its role in GSCs, was found to be critical for the survival of the cap cells. The autophagy defective cap cells are compromised in function and thereby unable to maintain GSCs during aging. We confirmed that deterioration of the GSC-niche, because of disrupted autophagy, not only results in apoptosis of cap cells in old age but also causes functional deterioration as measured by reduced niche-GSC/BMP signaling, which was observable from mid-life onwards.

Autophagy in other stem cells has been found to be important for their function or differentiation. Atg5 has been found to be important for self-renewal or differentiation of many adult stem cell types [42]. Based on numerous reports demonstrating the importance of autophagy for stem cell maintenance, we expected autophagy to be important for GSC maintenance as well. But surprisingly, in the case of female Drosophila GSCs, autophagy has been found to be dispensable. Previously, we suggested that GSCs may be protected from starvation-induced autophagy [34]. In this study, our approach for measuring basal autophagy flux in GSCs and assessing GSC maintenance has the following advantages. The data was quantified by sampling the entire volumetric fraction of GSCs. Conditions for inhibition as well as elevation of autophagy, namely chloroquine treatment and overexpression of autophagy gene (*Atg5*) as prescribed by the guidelines to assess autophagy, were included [4]. Disruption of autophagy in *Atg5* mutant GSCs was demonstrated. The study by Zhao and colleagues corroborates these findings; they also reported that autophagy in GSCs is low and that GSC maintenance is unaffected in *Atg6* and *Fip200* mutant GSCs [43]. Also, Sênos Demarco and colleagues reported that knocking down multiple core autophagy genes (*Atg1, Atg5, Atg6, Atg7, Atg8a/8b*) in male GSCs did not affect their numbers at ten days post-eclosion [18]. Taken together, these data suggest that in male and female GSCs, autophagy is dispensable.

How autophagy is regulated within GSCs is an interesting question, as autophagy levels are moderate to high in the neighboring cysts. Zhao and colleagues speculated that the reason for low autophagy in the GSCs was associated with the activation of BMP signaling. This speculation could be supported by the study published by Varga and colleagues [44], who demonstrated that BMP inhibits autophagy in male GSCs in Drosophila. Since the BMP-pMad signaling axis maintains female GSCs, these data strongly suggest that the GSC self-renewal program inhibits autophagy in the female GSCs.

Since higher autophagy is observed in cysts, it is likely that the differentiating factors upregulate autophagy, which is supported by Varga et al. 2022. Further, defective mitochondria are cleared by mitophagy in the female germline and are regulated by Atg1 [17,45]. Insulin/InR-mTOR signaling as been reported to control GSC division and growth and it could regulate autophagy alone or in combination with BMP signaling.

Autophagy in GSCs appears to be under tight control as eGFP-Atg5 puncta rarely formed in the GSCs but were present in the differentiating cells when eGFP-Atg5 was overexpressed (Figure S1B). eGFP-Atg5 puncta are known to be localized to phagophore assembly sites, isolation membranes, and immature autophagosomes, therefore, this observation also corroborates that autophagy in GSCs is low. Alternatively, this could be due to a lack of other core autophagy proteins required to form these intermediates. Similarly, Atg8a overexpression did not increase mCherry-Atg8a puncta formation, representing both autophagosomes and autolysosomes (Murmu and Shravage, in preparation). In contrast, previous studies from the murine model have shown that ubiquitous Atg5 overexpression and neuronal Atg8a overexpression in Drosophila are sufficient to elevate autophagy [9,39]. Thus, autophagy in GSCs may be tightly and differently regulated as compared to the differentiating germline cells.

Autophagy has also been shown to be important for stem cell maintenance during aging, particularly a direct role has been shown for hematopoietic and muscle stem cells [15,16]. However, a few studies demonstrate that *Atg5* deletion did not affect stem cell maintenance. For example, *Atg12* and *Atg5* conditional knockout HSCs in mice were maintained at normal numbers over time and *Atg5* deficiency did not affect hematopoietic stem cell maintenance in mice [16]. In case of murine neural stem cells, deletion for *Atg5*, *Atg16L1*, *Atg7* or *FIP200* all showed defective autophagy and accumulation of mitochondria, but only *FIP200* deletion affected NSCs maintenance [10]. In mice, deletion of *Atg5* did not affect neural stem cell or hematopoietic stem cell maintenance. Why Atg5 is required for autophagy in these stem cells but not for stem cell maintenance is an open question.

Sênos Demarco and colleagues showed that, for male GSC maintenance, autophagy is required non-autonomously in the supporting somatic CySCs that are part of the male GSC-niche [18]. Our observations are similar for female GSCs and their niche i.e., cap cells. The dichotomy between soma and germline makes the ovarian tissue interesting. Autophagy is regulated differently in these cells and has distinct functions. For instance, Barth and colleagues found that Atg1 and Atg13 were required in the soma i.e. follicle cells, but were dispensable in the germ cells for egg development [37]. We observe a similar phenomenon in the GSC-niche; autophagy is required in the somatic cells; cap cells and is dispensable in GSCs. Follicle stem cells are of somatic origin and are highly influenced by autophagy. Autophagy in follicle stem cells increases with age and also leads to their loss [46]. Recent studies have demonstrated that distinct cell types exist within the germarium and have different transcription profiles [47]. The germ cell transcriptional signature matched prominent features of the germline, including autophagy. In summary, autophagy has distinct functions in the female Drosophila germline and GSCs.

Cap cells are terminally differentiated post-mitotic cells. The data from Pan and colleagues shows that the cap cell population remains more or less unchanged throughout the lifespan of the flies [31]. Significant cap cell loss does not occur until seven weeks; even so the loss is less than 20%. Autophagy has been found to be important for the longevity of terminally differentiated post-mitotic cells like neurons. Despite the disruption of autophagy throughout the lifespan, we observed cap cell death only 40 days onwards. This could be due to RNAi-mediated knockdown not completely eliminating Atg5, although severe accumulation of Ref(2)P was observed. Several groups have described the role of apoptosis and autophagy in cell death during oogenesis [35,36,48]. It appears that both these contribute to cell death in cell type and context dependent manner. During early oogenesis autophagy was shown to be upstream of DNA fragmentation during apoptosis [36]. During mid-oogenesis cell death is driven majorly by apoptosis. At later stages during nurse cell death autophagy is the primary process that mediates cell death. In cap cell death, autophagy appears to be upstream of apoptosis. At the same time, although cell death was observed from 40 days onwards, deterioration of cap cell function was observed from 30 days onwards, suggesting disruption of cellular homeostasis. Since cap cells are not mitotically active in the adult ovary, mosaic analysis in these cells is challenging. However, such studies could be conducted by inducing *Atg* mutant clones during cap cell specification and division.

Although there is a significant and thorough understanding of the role of autophagy in stem cells as an intrinsic factor, there are very few reports connecting autophagy—niche—stem cells. A recent study has demonstrated that sensory nerves, that are part of the mesenchymal stem cell niche, activate autophagy in the mesenchymal stem cells via FGF1 (fibroblast growth factor 1), which is essential for mesenchymal stem cell maintenance [49]. A recent review consolidates literature from the point of view that the systemic nature of autophagy can be implicated to have a protective role in the maintenance of vascular niche homeostasis[50].

Our finding that GSC maintenance is autophagy-independent is baffling, especially considering the nature of these cells. The first level of distinction is that they are stem cells, and uniquely among the type of stem cells, they are GSCs. This is the first study that describes a novel role of autophagy in the niche cells for stem cell maintenance. This study exemplifies a model to study the effect of autophagy-defective niches on the stem cells that they harbor. Atg5 is conserved in metazoans and plays an important role in autophagosome formation and closure. Thus, these findings could have critical implications for understanding the role of autophagy in niche regulated stem cells including in diseases such as cancer. Our findings add to the understanding of the fundamental role of autophagy in stem cell maintenance and where it has a non-cell-autonomous effect.

## Materials and methods

### Fly maintenance

Flies were maintained at standard conditions; 25℃, 60-70% relative humidity, 12-12-hour light-dark cycle. Fly food composition per liter of food; 80 g sugar, 75 g corn flour, 30 g yeast, 30 g malt extract, 10 g agar, 0.12% methyl benzoate, 0.4% propionic acid and 0.08% orthophosphoric acid. For setting crosses, 0–5-day old males and virgin females were used. Until dissection, the flies of desired genotypes were housed as 15 females and 7-10 males in a vial which were flipped every two to three days to fresh food vials supplemented with dry yeast pellets. Two days prior to dissection, the ovaries were fattened by transferring the flies each day on fresh food vials supplemented with dry yeast pellets. Diethyl ether vapors were used to anesthetize flies. For chloroquine treatment, a stock solution of 50 mg/ml chloroquine in water was added to fly food to a final concentration of 3 mg/ml when the food cooled down to 50-60℃ during preparation. Flies of all experimental conditions were first fed on food supplemented with yeast pellets for two days. The flies for chloroquine treatment were transferred to fresh chloroquine containing food vials for two days while the control flies were transferred to fresh vials containing only normal fly food.

### Aging

Large crosses were set to collect a large number of age-synchronized progeny of the desired genotype which were collected within 24—48 hours of eclosion. Until dissection the flies were housed under standard conditions as 15 females and 7-10 males in a vial which were flipped every three to four days to fresh food vials supplemented with dry yeast pellets. The vials were kept horizontal throughout the duration in order to avoid death of flies due to sticking onto the food surface. Subsets of the collected and aged flies were dissected at each of the timepoints.

### Recombination of Atg5 mutant and FRT19A and its validation

The *Atg5^5cc5^* mutant was first crossed to isoFRT19A and the virgin female progeny was crossed to FM7i males. After a day of setting the cross, the flies were transferred to vials containing 500 μg/ml geneticin (G418)(Gibco, 11811-023) for egg laying. G418 was used to screen the FRT19A positive recombinants since the FRT19A cassette has neomycin as a selection marker. All the surviving virgin female progeny developed from the G418 vials were crossed to FM7i males in multiple single pair crosses. Males from the progeny of each of the single pair crosses were used to perform single fly PCR to confirm the presence of mutation using primers that specifically bind in the region of *Atg5* which was deleted in *Atg5^5cc5^* (forward primer: 5’GCACTACATGTCCTGCCTGA, reverse primer: 5’AGATTCGCAGGGGAATGTTT).

### Genotypes of all fly stocks

*+; +; + (OregonR), w118; +; +, w; +; nosGal4VP16 (BL-4937), w; nosP-mCherry-Atg8a; +, w; nosP-mCherry-Atg8a/CyO; nosGal4VP16/TM6b, Hu, y w; +; UASp-eGFP-drAtg5 (derived from BL-59848), y sc* v; Atg1 RNAi; + (BL-44034), y sc* v; +; Atg5 RNAi (BL-34899), y sc* v; Atg8a RNAi; + (BL-58309), y sc* v; +; Atg9 RNAi (BL-34901), y sc* v; Atg13 RNAi; + (BL-40861), y sc* v; +; Atg16 RNAi (BL-34358), w; If/CyO; HhGal4, UAS-GFP/TM6B, Tb, w; +; HhGal4, UAS-GFP/TM6B, Tb, w; +; Bab1Gal4/TM6B (BL-6803), w; TjGal4/CyO; +, y w; 3xmCherry-Atg8a; +, w; 3xmCherry-Atg8a/CyO; HhGal4, UAS-GFP/TM6B, y w^+^ Atg5^5cc5^ FRT19A/ FM7i; +; +, y w iso FRT19A; +; +, w hsFLP12, Ubi-RFP FRT19A; +; + (BL-31418), y w hsFLP12, His2Av.GFP FRT19A/FM7a; +; + (BL-32045)*.

### FLP-FRT based GSC clone generation and GSC retention assay

Large cross of *Atg5^5cc5^ FRT19A* with *whsFLP, Ubi-RFP FRT19A* for Atg5 mutant GSC clones and another cross of *isoFRT19A* with *whsFLP, Ubi-RFP FRT19A* for control GSC clones were set in multiple vials. The crosses were transferred to a set of fresh vials supplemented with yeast paste and removed after six-seven hours to obtain synchronized egg lay. The vials were subjected to heat shock for four consecutive days on seventh through tenth day after egg lay i.e., during pupal development. Heat shocks were applied for 50 minutes twice a day, six-eight hours apart, at 37℃ in a water bath. Only the flies which eclosed after the complete heat shock regime were collected. Therefore, the days-post-eclosion is the same as days-post-heat-shock. Females of the appropriate genotype were collected and aged until the stipulated timepoints. For each time-point, 8-10 of the collected females were dissected two, nine and nineteen days-post-heat-shock and immunostained for hts (1B1) and Lamin C for identification and quantification of RFP^+^ and RFP^−^ GSCs.

Immunostaining of Ref(2)P in Atg5 mutant GSCs was performed in two experiments. First using *Ubi-RFP FRT19A* where the progeny was heat shocked in the adult stage ten days post eclosion for three consecutive days in the regime stated above, and dissections were performed two days after the last heat shock. Second where the immunostaining was performed two-days post eclosion after pupal heat shock. The same procedure was repeated while using *His2AvGFP FRT19A* and the phenotype could be replicated (data not shown).

### Immunostaining

Flies were briefly anesthetized and dissected in Grace’s medium (Gibco, 11667-037). Ovaries were extracted and non-vitellogenic region of the ovarioles were partially teased apart using minutien pins. The ovaries were then transferred to 0.5 ml tubes with the aid of a cut-tip passivated with bovine serum albumin (BSA)(MP Biomedicals, 199897). The ovaries were fixed with 350 μl 4% paraformaldehyde (Sigma Aldrich, P6148) in 1xPBS (phosphate-buffered saline) pH 7.4 for 15 minutes at room temperature with the gentle nutation. All the following steps are performed with gentle nutation 15-20 RPM. The fixative was washed off with three washes of 400 μl 0.1% PBTx (triton-X-100{SRL, 64518} in 1xPBS) for five minutes each. Blocking was performed in 300 μl of 0.5% BSA in 1% PBTx for one hour at room temperature. The sample was incubated with at least 100 μl of the appropriate primary antibody solution in which the antibody was diluted in 0.3% PBTx containing 0.5% BSA. The incubation with primary antibody solution was performed at 4℃ overnight with gentle nutation 5 RPM. The primary antibody was washed off with 400 μl 0.1% PBTx for 15 minutes at room temperature. Following which the samples were again blocked for secondary antibody staining with 400 μl of 10% normal goat serum (NGS)(MP Biomedicals, 2939149) in 0.1% PBTx for two hours at room temperature. The secondary antibody diluted in the 10% NGS solution in 0.1% PBTx was incubated with the samples for two hours at room temperature. Three washes of 15 minutes each with 400 μl of 0.1% PBTx were performed at room temperature. For DAPI staining, the samples were incubated in 1 μg/ml DAPI in 0.1% PBTx solution for 10 minutes at room temperature and consequently washed off twice for five minutes each with 400 μl 0.1% PBTx. The samples were stored at 4℃ until mounting. All the PBTx solution was removed carefully and mounting medium SlowFade Glass mountant (Invitrogen, S36917) was added. All the ovaries were transferred onto a slide along with the mounting medium. In order to obtain flat mounting of the germarium, the region of the ovarioles which have the string of pre-vitellogenic stages and germarium at the tip were separated using minutien pins and the remaining large part of the ovaries was picked and removed from the slide. For optimal Confocal microscopy, ∼170 μm thick no.1 coverslips were used. Nail varnish was used to seal the slide. The slides were stored at 4°C protected from light until microscopy.

The following primary antibodies were used with the mentioned dilutions; anti-Cathepsin L (Abcam, ab58991; 1:400), anti-Ref(2)P (Abcam, ab178440; 1:1000), anti-Hts (DSHB, 1B1; 1:50), anti-Lamin C (DSHB, LC28.26; 1:50), anti-pMad (Abcam, ab52903; 1:50). The following secondary antibodies were used at 1:250 dilution; Goat anti-Rabbit Alexa Fluor 555 (Thermo Fisher Scientific, A21429), Goat anti-Rabbit Alexa Fluor 647 (Thermo Fisher Scientific, A21245) and Goat anti-Mouse Alexa Fluor 647 (Thermo Fisher Scientific, A21236).

### TUNEL staining

In Situ Cell Death Detection Kit, TMR red (Roche, 12156792910) was used for TUNEL staining. The procedure was repeated exactly the same for all five time points. Ovaries were dissected and teased as described for immunostaining. All the following steps were performed with gentle shaking/neuation and the volumes indicated are for each sample. Fixation: 15 minutes at room temperature, 300 μl 4% paraformaldehyde in PBS pH 7.4. Followed by three washes of 5 minutes each at room temperature with 300 μl 0.1% PBTx for each. Subsequently, blocking and permeabilization: 300 μl of 1% PBTx containing 0.5% BSA, for one hour at room temperature. Followed by one wash with 400 μl 1x PBS for 2 minutes at room temperature. Total of 250 μl TUNEL reaction solution was prepared during each experiment, which comprised of 225 μl TUNEL TMR label solution and 25 μl TdT enzyme solution. 50 μl TUNEL reaction solution was used per sample. TUNEL reaction was performed by adding the TUNEL reaction solution to the sample and incubating for 2 hours at 37℃ in a shaking incubator, 80 RPM and protected from light. All following steps were performed with samples protected from light. The samples were proceeded for immunostaining after blocking in 300 μl of 1% PBTx containing 0.5% BSA for 30 minutes at room temperature.

### Confocal microscopy

The imaging was performed on the Leica SP8 Confocal microscope or Confocal mode of the Zeiss LSM900 Airyscan microscope. All the microscopy performed on the Leica SP8 was done using 63x oil immersion objective NA=1.4. For all intensity and puncta count analyses performed on Leica SP8, the settings were as follows; resolution=1024×1024 pixels, scan speed=100 Hz, bit depth=8-bit. For obtaining z-stacks for GSC or cap cell counting; resolution=512×512 pixels, scan speed=400-600 Hz, bit depth=8-bit, z-step size=0.5 μm.

For the measurement of autophagy flux in GSCs, z-stacks spanning entire GSCs were acquired by visual confirmation from the DAPI (nucleus) channel. The z-step size was set to 0.4 μm which ensured that puncta were not missed out between steps. 63x oil immersion objective NA=1.4, 512×512 pixels at 0.05 μm per pixel resolution, scan speed=8, bit depth=16-bit.

For all GSC and cap cell counts, z-stacks of the tip of the germarium which covered the entire GSC niche were acquired by visual confirmation (z-step size=0.5 μm). While imaging for Ref(2)P puncta in the cap cells, the central plane of the cap cells was focused using the Lamin C channel. While imaging 3xmCherry-Atg8a in cap cells, the plane maximally covering the cluster of cap cells centrally was identified in the DAPI channel. The GSCs were identified by their location and size of the nucleus and the optical section across the central plane was acquired by focusing in the DAPI channel for the following experiments; Ref(2)P in Atg5 knockdown, Atg5 overexpression, Atg5 mutant, and pMad. Imaging settings for all intensity and puncta count experiments were exactly the same for all experimental conditions and the imaging of a set was completed in one imaging session.

### Image analysis

All the image analyses involving intensity measurement or puncta count were performed in ImageJ and Fiji. All steps performed for image analyses were exactly the same for all the control and test images in a data set. For GSC counts, Atg5 mutant and control GSC clone counts, cap cell counts and TUNEL positive cap cell counts, LAS-X and Zeiss Zen 3.7 software were used to visualize and manually count the cells. GSCs were unequivocally identified by the following characteristics; their location, size of the nucleus, presence and orientation of the spectrosome. Cap cells were identified by their nuclear shape, location and Lamin C ring.

For autophagic flux determination in the GSCs, nosP-mCherry-Atg8a and Cathepsin L puncta within the GSCs were manually marked in the plane with maximal area and intensity. Additionally, colocalized mCherry-Atg8a and Cathepsin L puncta were also manually identified. Similarly, 3xmCherry-Atg8a puncta in the cap cells were manually quantified. For quantification of Ref(2)P puncta in the GSCs or cap cells; ROI was drawn to define either the GSCs or cap cells in the DAPI channel and the ROI was imported into the ‘ROI manager’. The Ref(2)P channel was first subjected to the ‘Subtract background’ function with rolling ball radius 50 pixels or 35 pixels according to the optimal value for the data set. The resultant images were thresholded referring to the display under ‘Max entropy’ algorithm and then manually adjusted for each image such that the background was excluded but the true signal of the puncta did not erode. The thresholded images were then used for the function ‘Analyze particles’ in which the following parameters were set; Size (micron^2): 0.02-infinity, Circularity: 0-1, from which the output was obtained as number and size of puncta within the marked ROI. pMad intensity in the GSCs was quantified in ImageJ by marking ROIs and recording the ‘mean intensity’.

### Statistical analysis

For all comparative analyses, Student’s t-test assuming unequal variance was used. For statistical comparison of the proportion of TUNEL positive cap cells out of total cap cells among different genotypes, z-test for two proportions was used. Microsoft Excel was used to store, arrange and analyze data. Graph Pad Prism was used for plotting all the graphs.

## Data sharing

All data are available from the authors upon request.

## Acknowledgments

We thank Dr. Gábor Juhász for providing the Atg5 mutant and 3xmCherry-Atg8a fly stocks, Dr. Manish Jaiswal for providing the FRT19A stocks and Dr. Richa Rikhy and the IISER-Pune fly facility for providing many essential fly stocks. We thank ARI Confocal facility for help with imaging. We thank members of the Shravage lab for helpful discussions. We thank Dr. P. K Dhakephalkar, Director, Agharkar Research Institute, Pune, India, and the Developmental Biology fraternity for support and facility access. This work was supported by DST-SERB grant number ECR/2015/000239, BT/RLF/Re-entry/58/2013 and BT/PR12718/MED/31/298/2015 to BVS. KN was supported by ICMR-SRF 2020-6879/CMB-BMS and is a registered Ph.D. student affiliated to Department of Biotechnology, Savitribai Phule Pune University (SPPU), Pune, India (Registration No. 175, PGS/4204). BVS is affiliated to Savitribai Phule Pune University (SPPU), Pune, India, and is recognized by SPPU as PhD guide in Biotechnology and Zoology. We apologize for not being able to cite research work in the field owing to space limitations.

---

**Supplementary Figure S1:**
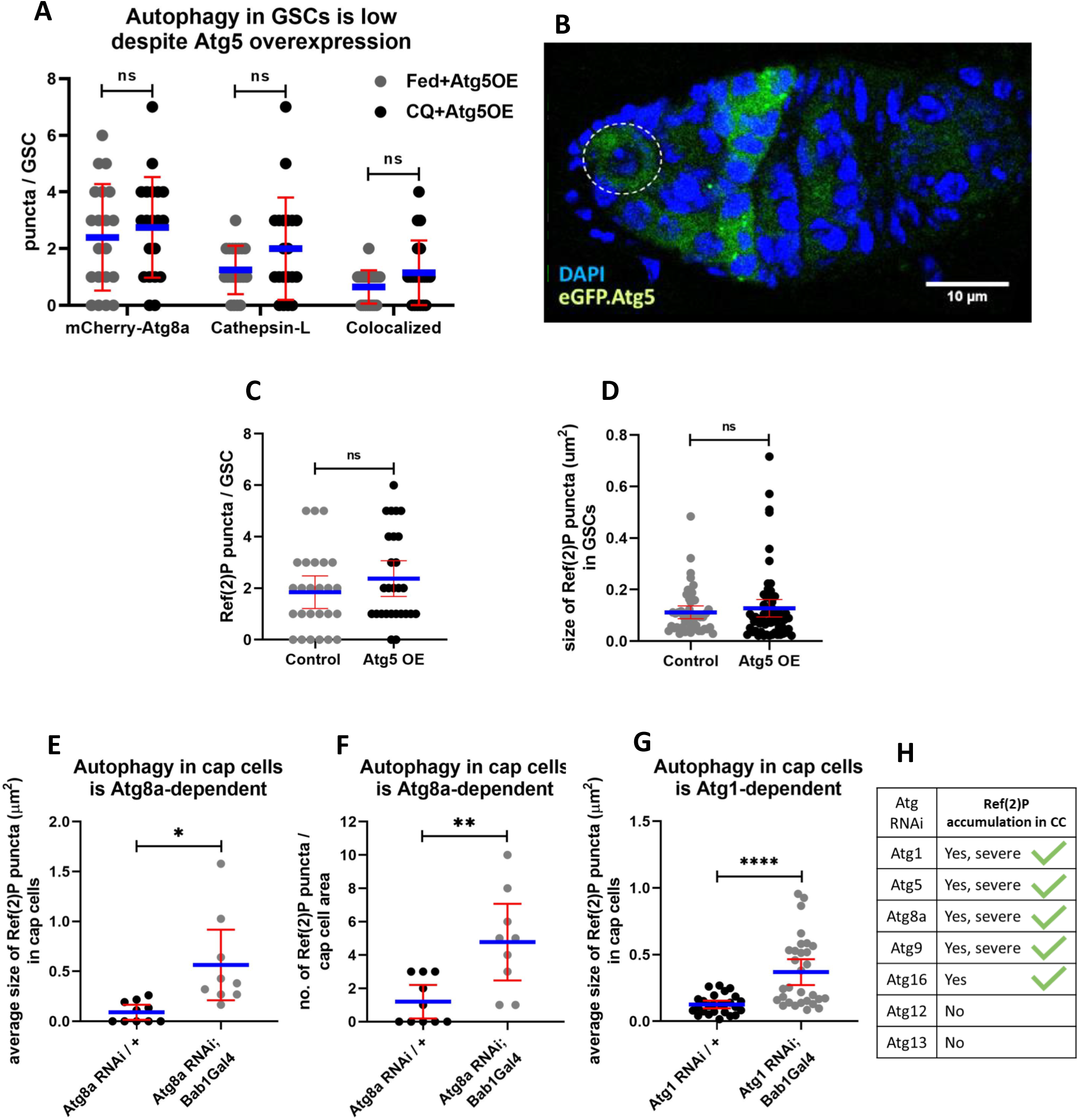
(A) Interleaved scatter plot showing no difference in number of autophagic vesicles in Atg5 overexpression (Atg5 OE) GSCs upon CQ treatment. Blue line represents the average and error bars represent standard deviation. n=20 GSCs per treatment. (B) Representative image of a cross-section of the central plane of the germarium showing eGFP fluorescence from eGFP-Atg5 (Atg5 overexpression). eGFP fluorescence is visible in the GSC which marked by the dashed outline and bright puncta are visible in region 2. Interleaved scatter plots showing no change in the (C) number and (D) size of Ref(2)P puncta in GSCs upon Atg5 overexpression. n=27±1 GSCs per genotype. Interleaved scatter plots showing increase in size (E) and number (F) of Ref(2)P puncta upon *Atg8a* knockdown in the cap cells and the same for size of puncta upon *Atg1* knockdown (G) in the cap cells. Blue line represents the average and error bars represent standard deviation. n=10±1 (Atg8a KD) and n=31±1 (Atg1 KD) cap cell planes/area per genotype. (H) List of all the Atgs tested for knockdown in niche and the observation of Ref(2)P accumulation in cap cells \**p*<0.05, \*\**p*<0.01, \*\*\*\**p*<0.0001.

**Supplementary Figure S2:**
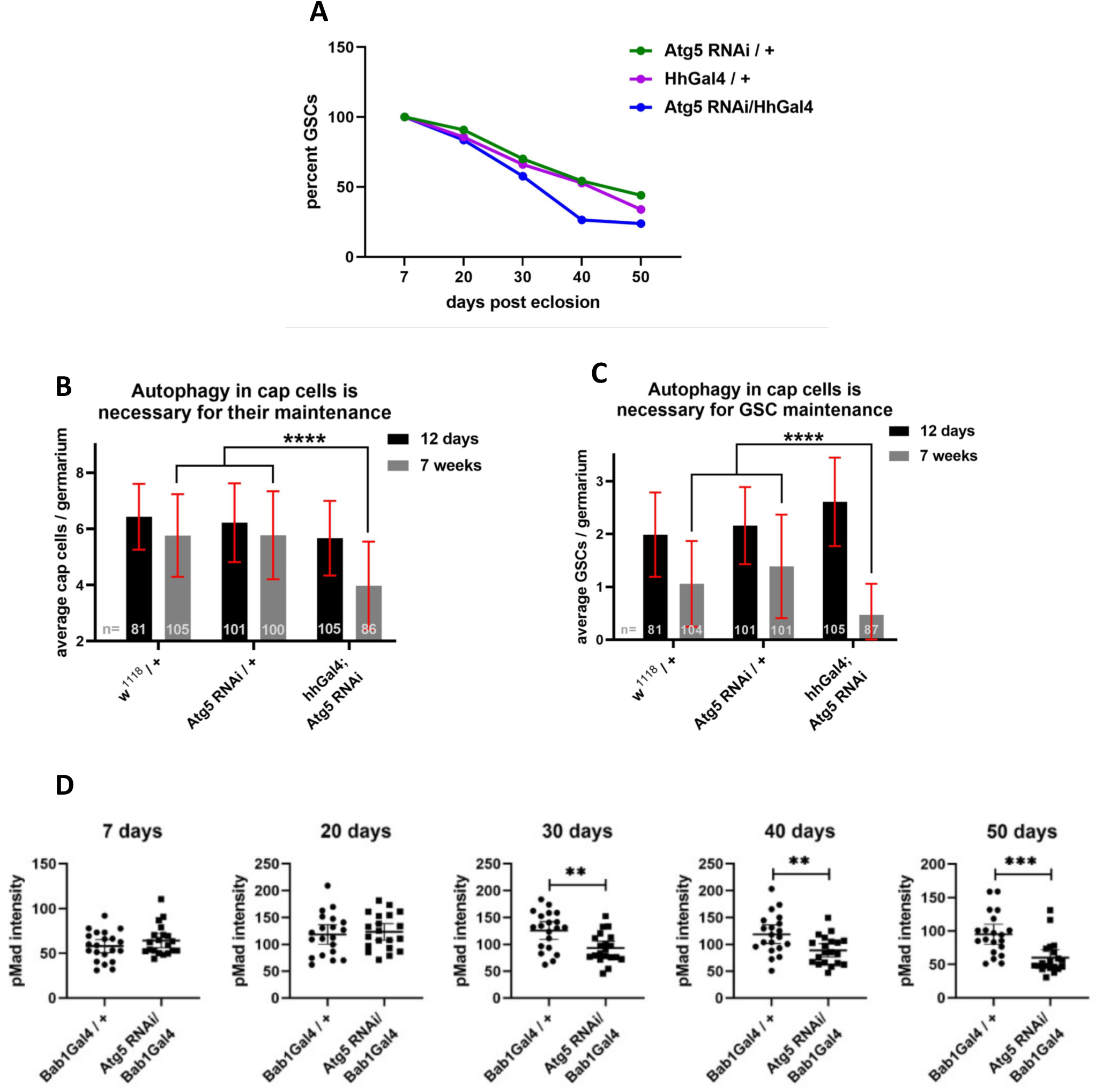
(A) Line graph showing rapid loss of GSCs during aging due to *Atg5* knockdown in cap cells. Bar graphs showing that the initial number of cells at 12 days is comparable among all genotypes for cap cells (B) and GSCs (B) and the exacerbated loss of both the cell types during aging is attributable to disrupted autophagy in cap cells due to *Atg5* knockdown. Error bars represent standard deviation. Sample sizes are the number of germaria as indicated in the graphs. (D) Interleaved scatter plots showing pMad intensity in GSCs of the mentioned genotypes at the five mentioned timepoints. Error bars represent standard deviation. n=20±2 GSCs per genotype per time point. \*\**p*<0.01, \*\*\**p*<0.001.

## Abbreviations

*Atg5*/Atg5: Autophagy-related gene/protein 5
GSC: Germline Stem cell
BMP: Bone Morphogenetic Protein
*Dpp*: Decapentaplegic
*Mad*: Mothers against Dpp
pMad: phospho-Mad
*Ref(2)P*: Refractory to Sigma P
*nosP*: nanos promoter (driven)
*FLP-FRT*: Flippase-Flippase Recombination Target

